# Additional analyses exploring the hypothesized transdifferentiation of plasmablasts to developing neutrophils in severe COVID-19

**DOI:** 10.1101/2020.10.15.339473

**Authors:** Aaron J. Wilk, Arjun Rustagi, Nancy Q. Zhao, Beth A. Martin, Angela J. Rogers, Catherine A. Blish

## Abstract

We thank Alquicira-Hernandez et al. for their reanalysis of our single-cell transcriptomic dataset profiling peripheral immune responses to severe COVID-19. We agree that careful analysis of single-cell sequencing data is important for generating cogent hypotheses but find several aspects of their criticism of our analysis to be problematic. Here we respond briefly to misunderstandings and inaccuracies in their commentary that may have led to misinformed interpretation of our results.

## Main

Alquicira-Hernandez et al.^1^ question the plausibility of the potential lineage relationships between plasmablasts and developing neutrophils that we postulated as a part of our recent work^2^. We appreciate their commentary and concur that careful computational analysis of single-cell RNA sequencing (scRNA-seq) data is necessary. Our study, the first to publicly share scRNA-seq data to profile immunity in COVID-19, was by its design and execution descriptive, correlative, and hypothesis-generating, given the limitations of the dataset acknowledged in our original manuscript. Our goal was to develop a resource for the scientific community to better understand COVID-19, and to identify distinctive immune features for further study. We regret that we may have not adequately conveyed the hypothesis-generating nature of our study; if any reader came away with the impression that we had claimed to have “proven” the existence of a plasmablast to neutrophil transition, this was not our intent, and we apologize.

We chose our words very carefully when describing our findings, explicitly choosing not to say that we had “proved” or “concluded” new hypotheses with scRNA-seq data alone, particularly as it related to a potential transdifferentiation pathway. In this regard, we point to our original manuscript rather than Alquicira-Hernandez et al.’s paraphrasing, which left out important context for our stated conclusions. To wit, our final statement on this putative pathway reads, “Collectively, we observe a developing neutrophil population that may be characteristic of ARDS in severe COVID-19 infection; our data suggest that these cells may derive from plasmablasts, but they may also represent developing neutrophils derived from emergency granulopoiesis”^2^.

Alquicira-Hernandez et al. argue that transdifferentiation between plasmablasts and developing neutrophils is biologically implausible and, therefore, that the association between these two cell types in uniform manifold approximation and projection (UMAP) space must represent an artifact of the computational pipeline we selected. Alquicira-Hernandez et al. thus re-analyzed our data with different preprocessing recipes to see if the phenotypic association between plasmablasts and developing neutrophils would break, implying that the relationship between these cells would be artifactual.

Our response to this argument is, as follows:

1. Alquicira-Hernandez et al. assert that we drew our hypothesis solely from the proximity of plasmablasts and developing neutrophils in non-linear dimensionality reduction space, which is incorrect. Our hypothesis was based primarily on a cellular trajectory analysis by RNA velocity (Figure 4 of our original manuscript). This orthogonal computational technique uses the kinetics of RNA splicing to calculate a time derivative of gene expression, which can computationally infer the trajectory of cellular differentiation^3^,^4^. Based on this analysis, we believe our hypothesis is plausible because we observed sequential downregulation of genes associated with plasmablasts and upregulation of genes associated with neutrophil development across the inferred latent time trajectory. This was coincident with upregulation of C/EBP transcription factors known to drive neutrophil development, and consistent with the pattern observed previously during B cell to macrophage transdifferentiation^5^–^9^. Thus, the results of RNA velocity-based analysis led us to postulate that the phenotypic relationship between plasmablasts and developing neutrophils could represent transition between the two cell types.
2. Alquicira-Hernandez et al.’s argument takes two inherently contradictory positions on the sufficiency of our dataset to prove, or disprove, a plasmablast-to-neutrophil transition in COVID-19. Given that we have stated that our data is insufficient to make such a firm conclusion, we do not find that Alquicira-Hernandez et al. could definitively disprove this putative transition using the same data alone.
3. This argument overextends the interpretability of scRNA-seq data by overemphasizing the role of parameter tuning in preprocessing. While we agree that careful selection of preprocessing parameters is an essential component of scRNA-seq data analysis, there are nonetheless a plethora of reasonable ways to analyze and visualize a scRNA-seq dataset. For many of these parameters, there is no widely-accepted formal method of determining what is the “best” to use for each dataset. For example, Alquicira-Hernandez et al. argue that we may have overfit our data by regressing the number of unique molecular identifiers (UMIs) and genes detected per cell, but the impact of such potential overfitting is likely to be inconsequential given the extremely high ratio of variables to covariates.

To explore this last idea further and extend the reanalysis performed by Alquicira-Hernandez et al., we have conducted a second analysis of our dataset using 24 different combinations of covariates and two other parameters not discussed by Alquicira-Hernandez et al. These additional parameters are the number of highly-variable genes (nHVG) used for transformation and dimensionality reduction, and the number of principal components (nPC) used for dimensionality reduction. This second analysis reveals that, in the vast majority of combinations, there is still a phenotypic association between plasmablasts and developing neutrophils (Figure 1a). This is similar to the UMAP projections generated by Alquicira-Hernandez et al., which still show developing neutrophils closely associated with plasmablasts, with several plasmablasts embedding closer to developing neutrophils than with other plasmablasts (Figure 1a, purple box). To examine this phenotypic relationship outside of the non-linear dimensionality reduction manifold space of UMAP, we additionally hierarchically clustered pseudobulk average gene expression profiles of each cell type in our dataset, which again indicates that plasmablasts and developing neutrophils are phenotypically related (Figure 1b). It is important to note that this does not indicate relationship in terms of cell lineage, but merely relationship of transcriptional phenotype. Taken together, these analyses re-affirm our previous decision to explore this relationship further with RNA velocity analysis.

**Figure 1.**
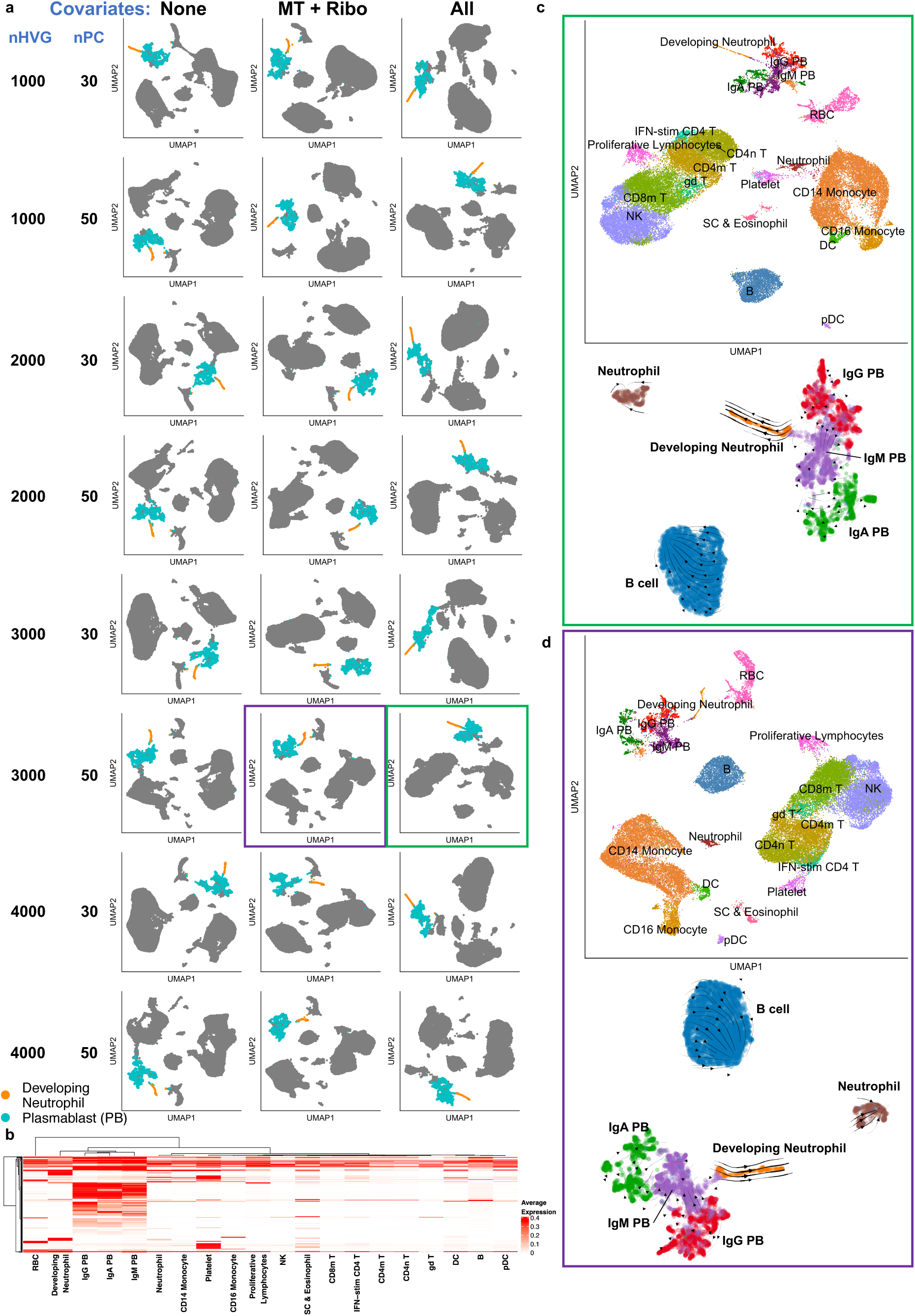
scRNA-seq data processed with different preprocessing parameters supports the hypothesis of plasmablast-developing neutrophil transdifferentiation. **a)** UMAP embeddings generated with 24 different combinations of regressed covariates, number of highly-variable genes (nHVG), and number of principal components for dimensionality reduction (nPC). “None”, no covariates regressed; “MT + Ribo”, only percentage of mitochondrial and ribosomal reads regressed; “All”, percentage of mitochondrial and ribosomal reads, as well as number of UMIs and number of genes detected per cell regressed. Only developing neutrophils and plasmablasts are colored; all other cell types are gray. “PB”, plasmablast. **b)** Hierarchically-clustered pseudobulk average expression profiles of the top 250 HVG for each cell type. **c)** UMAP dimensionality reduction projection of full dataset generated using original preprocessing parameters colored by cell type (top; green box corresponds to embedding shown in panel (**a**)). UMAP embedding of plasmablasts, developing neutrophils, B cells, and mature neutrophils generated using original preprocessing parameters overlaid with RNA velocity stream (bottom). **d)** Same plots as described in (**c**), using preprocessing parameters used by Alquicira-Hernandez et al. (percentage of mitochondrial and ribosomal reads regressed, 3000 highly-variable genes, 50 principal components).

We performed RNA velocity analysis using preprocessing parameters employed in our original manuscript (Figure 1c, green box) and using preprocessing parameters used by Alquicira-Hernandez et al. (Figure 1d, purple box). To analyze the dynamic relationship between additional cell types that may be biologically related to plasmablasts and developing neutrophils, we embedded only plasmablasts, developing neutrophils, B cells, and a population of low-density mature neutrophils we identified. We found that, with both sets of preprocessing parameters, developing neutrophils appear to transition from plasmablasts and do not occupy similar UMAP manifold space as B cells and mature neutrophils (Figure 1c, d).

Alquicira-Hernandez et al. hypothesized that plasmablasts should be more closely related to B cells than developing neutrophils, and that developing neutrophils should be phenotypically associated with mature neutrophils. Upon finding that an embedding of these four cell types alone showed a relative lack in relatedness between plasmablasts and B cells, the authors concluded that plasmablasts and developing neutrophils must be misclassified as related cell types. However, it is incorrect to assume B cells and plasmablasts should be phenotypically related in UMAP space, as these cell types are dramatically different in terms of gene module expression (eg. proliferation) that is easily detected at the transcriptional level (Figure 1b), and because the kinetics of B cell-to-plasmablast differentiation in these patients may not enable identification of intermediate cell states in the periphery. While it does remain possible that developing neutrophils and plasmablasts are related in UMAP space because they are both proliferative cell types, there are other proliferative T and NK cells in the dataset that are not phenotypically related and this argument does not have bearing on trajectories predicted by RNA velocity. We thus conclude that our selection of preprocessing parameters was reasonable and would have led to the same hypotheses had we chosen different parameters.

Alquicira-Hernandez et al. also imply that we did not fully consider the possibility that hemophagocytic lymphohistiocytosis (HLH) could have explained our findings because of the difficulty in making this diagnosis. First, we would like to reiterate that such an explanation would be expected to result in an increase in complexity (# genes detected per # UMI sequenced per cell) of developing neutrophils, which we did not observe (Extended Data Figure 9 of our original manuscript^2^). While it is possible that internalization of one cell into another, either by emperipolesis or hemophagocytosis, could confound our interpretation, these mechanisms of intact cell ingestions are exceptionally rare behaviors of both neutrophils and B lymphocytes at any differentiation stage, and none of the published series of peripheral smears in COVID-19 have revealed this phenomenon^10^–^12^. In addition, none of the patients in our cohort had clinical characteristics of HLH and none received granulocyte colony-stimulating factor (G-CSF) or granulocyte-macrophage colony-stimulating factor (GM-CSF) therapy on the original authors’ review. To probe the possibility of HLH in our cohort more deeply, we requested an independent review of all available clinical data (including demographics, ethnicity, laboratory and imaging data, physical examinations, and clinical treatment plans) from our cohort by a hematologist and expert in adult HLH (Dr. Beth A. Martin). Standard scoring (HLH-94, H score) and expert assessment were utilized. The available clinical data was sufficient to conclude that no patient had HLH. Of note, no patient met criteria for bone marrow biopsy or other tissue evaluation for the presence of hemophagocytosis.

In conclusion, we would like to thank Alquicira-Hernandez et al. for their commentary and reanalysis of our work. While it remains possible that the phenotypic association and predicted trajectory dynamics between plasmablasts and developing neutrophils is an incidental finding, the additional analysis presented here indicates this finding is not an artifact of our analytical pipeline. We believe that a plasmablast-to-neutrophil transdifferentiation in severe COVID-19 remains an intriguing and plausible hypothesis, one that we are working to validate through isolation of the correct cell population and DNA sequencing of the BCR loci to conclusively determine the developmental origins of these cells. Until we are able to generate such data, we would like to reiterate that our previous manuscript is exploratory, observational, and does not claim to have demonstrated the veracity of this transition. Finally, we would agree with the authors’ title with a slight modification, that there is “No **direct, mechanistic** evidence that plasmablasts transdifferentiate into developing neutrophils in severe COVID-19”: our study was not designed to find this evidence.

## CONTRIBUTIONS

AJW performed bioinformatic analyses; BAM performed independent review of clinical data; AJW, AR, NQZ, AJR, and CAB wrote the manuscript.

## DATA AVAILABILITY

All data analyzed in this manuscript are publicly available, and relevant accessions and web links are provided in the original publication.

